## Supplementary Material for "Semantics across the globe: A universal neurocognitive semantic structure adaptive to climate"

**The PDF file includes:**

Supplementary Text S1 to S6
Figures S1 to S9
Tables S1 to S7

### Supplementary Texts

#### S1. Principal component analyses results

In addition to inter-language correlation analyses, we conducted a Principal Component Analysis (PCA) on 204 concepts common across all 53 languages to further assess semantic commonalities. This complementary approach treats languages as features and semantic patterns as samples (Figure S2a), allowing us to extract the first principal component (PC1) as an indicator of cross-linguistic commonalities. The variance explained by PC1 reflects the degree of universality in semantic structure (Cole et al., 2014; Romney et al., 2000). While both PCA and inter-language correlation methods aim to quantify cross-linguistic similarities, they differ in their approach to handling between-concept variations and the number of concepts analyzed. The PCA utilized a subset of 204 words shared across all 53 languages, concatenating dimensional vectors for each concept to construct comprehensive semantic structures.

Results (Figure S2b, upper panel) showed that the neurocognitive-dimensional representation exhibits the highest degree of universality (44.31%), followed by local-distributed (36.38%), feature-based (34.45%), and global-distributed (34.16%) models. Compared to statistical random models, only the neurocognitive model was consistently positioned at the upper bounds of the distributions, signifying a relatively universal semantic structure ($P < 0.0001$ and $P = 0.01$, respectively; Figure S2b, lower panel).

For the neurocognitive semantic representation, the scree plot (Figure S2b, upper panel) showed a dominant first component, with subsequent components explaining substantially less variance (e.g., PC2 explains 4%). PC1 showed positive correlations with all 53 languages ($M = 0.66$, $SD = 0.13$), contrasting with PC2's mixed positive and negative loadings ($M = -0.02$, $SD = 0.20$). Spanish demonstrated the highest PC1 correlation ($r = 0.83$), while Korean exhibited the lowest ($r = 0.35$; full results in Figure S2c). The PC1 space of the neurocognitive structure matrix (Figure S2d) showed a general division between sensory-motor-related and social-cognitive-related dimensions. Notably, the "sound" dimension clustered with social-cognitive dimensions, while the "space" dimension aligned with sensory-motor dimensions. The matrix also visualizes cross-linguistic agreement in the representation of specific concepts, such as the strong association of "star" across “shape”, “color”, and “time” dimensions.

#### S2 Validation using pretrained embedding models from Wikipedia and the Open Subtitles.

To validate the findings from Study 1 (language computation study), we conducted additional analyses using text embedding data derived from a combination of Wikipedia and the Open Subtitles database (Van Paridon & Thompson, 2021). This validation study examined semantic representations of 1,016 NorthEuralex (NEL) concepts across 35 languages, analyzing commonalities and variations in different semantic representations.

<u>Language samples.</u> Language sample selection followed a two-step process: First, we identified 43 languages with translated word forms available in the word embedding models. Second, we excluded 8 languages that were missing over 25% vector representations for NEL concepts, ensuring

adequate shared concept coverage between language pairs. This process resulted in a final sample of 35 languages.

Commonalities on the neurocognitive semantic structure. We replicated the analysis of commonalities captured by neurocognitive semantic structures, constructing different theoretical models (neurocognitive, distributional [local and global], and featural) using semantic projection methods. Cross-language commonalities were compared using two complementary methods: inter-language correlation and principal component analysis.

a) *Inter-language correlation results*. As shown in Figure S4a (left panel), the neurocognitive-dimensional semantic representation demonstrated greater cross-language similarity compared to other representation types (mean Fisher-z-transformed r: $M_{\text{neurocognitive}} = 0.77$, SD = 0.15; $M_{\text{distributional (global)}} = 0.53$, SD = 0.13; $M_{\text{distributional (local)}} = 0.44$, SD = 0.09; $M_{\text{semantic feature}} = 0.43$, SD = 0.11; neurocognitive vs. distributional (global): Wilcoxon test V = 177309, $P < 1 \times 10^{-16}$; neurocognitive vs. distributed (local): Wilcoxon test V = 177310, $P < 1 \times 10^{-16}$; neurocognitive vs. semantic feature: Wilcoxon test V = 177310, P < 1 × 10-16).

*b) Principal component analysis results.* We conducted a PCA for 35 languages and 354 NEL concepts shared across all languages. The variance explained by the first component (PC1) indicated that neurocognitive-based representations of concepts are more universal than other semantic structures (Figure S4a, right panel). The neurocognitive-dimensional representation exhibited the highest degree of universality (65.52%), followed by local-distributed (55.99%), global-distributed (49.66%), and feature-based models (47.55%). For the neurocognitive semantic representation, the scree plot showed a dominant first component, with subsequent components explaining substantially less variance (e.g., PC2 explains 3.4%). PC1 showed positive correlations with all 35 languages (M = 0.81, SD = 0.07), contrasting with PC2's mixed positive and negative loadings (M = -0.001, SD = 0.19).

Variations associated with environmental variables. Within a sample of 17 languages (136 language pairs) where all four environmental variables were available, they collectively explained 64% of the semantic (neurocognitive) space variations (Spearman $\rho = 0.81$; $P < 1 \times 10^{-16}$; Figure S4b). We replicated the association patterns between climate and neurocognitive semantic variations using a linear regression model to assess the unique effect of each environmental factor. As indicated in Figure S4b, climate showed the strongest unique explanatory effects (climate: $\beta = 0.44$, 95% CI = [0.33, 0.58], $P = 1.39 \times 10^{-10}$). Culture and linguistic history also showed significant unique effects (culture: $\beta = 0.22$, 95% CI = [0.12, 0.33], $P = 4.64 \times 10^{-5}$; linguistic history: $\beta = 0.28$, 95% CI = [0.20, 0.35], $P = 7.86 \times 10^{-12}$). Geographic distance showed negative effects on semantic variations ($\beta = -0.18$, 95% CI = [-0.29, -0.06], P = 0.002). The effects of climate on specific neurocognitive dimensions were also replicated (12 of 13 semantic dimensions, FDR-corrected *q*s < 0.05; shape, q = 0.06; Figure S4b, right panel).

#### S3 Statistical methods for controlling non-independence

In our study, we addressed potential biases arising from non-independent sampling of languages and subjects, particularly the spatial autocorrelations between samples, when estimating the effects of environmental variables. A primary concern was the non-independence in language sampling. While we incorporated other environmental variables into the same regression models as the potential controls, we implemented additional measures to ensure the robustness of our findings, particularly regarding the effects of environmental variables such as climate.

*a) Linear mixed effects regression models*. To account for the non-independent sampling of languages, we employed linear mixed regression models with a crossed random-effects structure. This approach nested language pairs within language families. Following the method suggested by Chen et al. (2017), we "doubled" the data (with redundancy) to allow for fully crossed random effects, accounting for the symmetric nature of the inter-language/inter-subject matrix and the fact that each participant contributes twice in the model. Therefore, prior to statistical inference, we manually adjusted the degrees of freedom to N - k, where N represents the number of unique observations and k denotes the number of fixed effects in the model. This correction ensures appropriate statistical power and reduces the risk of Type I errors. All reported findings in this article utilize these corrected degrees of freedom. The results of linear mixed effects regression models are presented in the main text.

*b) Additional validation approaches*. To further validate the robustness of the climate-semantics relationship, we implemented several complementary analytical strategies:

1. Alternative random effect structures: We tested more complex random effect specifications to rigorously control for hierarchical non-independence relationships. The enhanced model with nested random effects can be formally expressed as:

$$Semantic\_Distance_{ij} = \beta_0 + \beta_1 Climate_{ij} + (1 \mid Family_i/Language_i) + (1 \mid Family_j/Language_j) + \varepsilon_{ij} \quad (1)$$

, where i and j index the language pairs, $\beta_0$ represents the intercept, $\beta_1$ represent the fixed effects for climate, (1|Family_i/Language_i) and (1|Family_j/Language_j) represent the nested random intercepts where languages are nested within their respective language pairs (Language_ij) and language families (Family_ij), and ε_ij represents the residual error term. This nested structure accounts for the hierarchical nature of the data, where language pairs belong to specific language families, thus more rigorously controlling for phylogenetic non-independence in the data.

These models consistently demonstrated significant climate effects across all three studies (Study 1 – Wikipedia + CC: $\beta = 0.26$, 95% CI [0.20, 0.31], $P = 1.21 \times 10^{-18}$; Study 1 – Wikipedia + Subs: $\beta = 0.44$, 95% CI [0.34, 0.54], $P = 1.23 \times 10^{-14}$; Study 2: $\beta = 0.14$, 95% CI [0.05, 0.24], $P = 0.004$; Study 3: $\beta = 0.12$, 95% CI [0.06, 0.19], $P = 0.0002$), confirming that our findings are not artifacts of model specification.

2. Individual-level analysis: For Studies 2 and 3, we performed analyses at the individual subject dyad level rather than the language level. The model with the nested random effect's structure at this finer-grained level can be formally expressed as:

$$Semantic_{Distance_{ij}} = \beta_0 + \beta_1 Climate_{ij} + (1 \mid Family_i/Language_i/Subject_i) + (1 \mid Family_j/Language_j/Subject_j) + \varepsilon_{ij} \quad (2)$$

, where i and j index the individual subject pairs, $\beta_0$ represents the intercept, $\beta_1$ represents the fixed effect for climate, (1|Family_i/Language_i/Subject_i) and (1|Family_j/Language_j/Subject_j) represent the three-level nested random effects structure where subjects are nested within languages, which are in turn nested within language families. This structure captures the complete hierarchical nature of our data by accounting for non-independence at the subject level (participants who speak the same language), the language level (languages that belong to the same family), and the language family level. These analyses showed significant climate effects (Study 2: β = 0.18, 95% CI [0.17, 0.18], $P < 1 \times 10^{-16}$; Study 3: β = 0.07, 95% CI [0.03, 0.11], P = 0.0003), confirming that our findings persist at multiple levels of analysis.

**S4 Validation of commonalities in Study 2 and Study 3**

To validate commonalties in neurocognitive dimensional structures shared among individuals in different language groups, we employed a variation comparative analysis framework on within-language and cross-language subject pairs using human rating data (Study 2) and neural activation data (Study 3).

Study 2: Validation of commonalities (human rating data)

Methods: We constructed an inter-subject correlation matrix using Pearson correlation coefficients between subjects' ratings on 207 concepts across 13 dimensions (2,691 ratings per subject). To examine the existence of a language-specific component, we conducted an independent Wilcoxon signed-rank test (two-sided) comparing rating similarities between within-language and cross-language subject pairs. We then applied the "mean correlation" method (Romney et al., 2000) to quantify commonalities and variations across languages. The variance of this matrix was decomposed into 3 distinct components: a universal component shared by individuals regardless of their spoken language, a language-specific component that represents the average of unique semantic knowledge shared by individuals within a particular language group, and a component that captures individual variance not accounted for by language-specific variations. The square root of the mean correlation of certain subject pairs captures the proportion of knowledge shared among a specific group. We thus performed the following calculations: (i) compute the square root of averaged Pearson correlation of cross-language subject pairs, as measurement of a shared proportion (denoted by $\text{Var}^2_{\text{language-shared}}$); (ii) compute the square root of averaged correlation of within-language subject pairs for each language (denoted by $\text{Var}^2_{\text{shared-within}}$), then subtract the shared proportion from sample-size-weighted mean of within-language knowledge, as measurement of a language-specific proportion ($\text{Var}^2_{\text{shared-within}} - \text{Var}^2_{\text{language-shared}}$, denoted by $\text{Var}^2_{\text{language-specific}}$); and (iii) compute the individual variability portion by subtracting the shared proportion and language-specific variations from one ($1 - \text{Var}^2_{\text{language-shared}} - \text{Var}^2_{\text{language-specific}}$, denoted by $\text{Var}^2_{\text{individual}}$).

Results: We investigated the existence of a language-specific component by comparing within-language similarities (N = 3,764 pairs) to cross-language similarities (N = 55,992 pairs). Results showed higher within-language similarities (M = 0.55) compared to cross-language similarities (M = 0.47; Wilcoxon test W = 71,836,161, $P < 1 \times 10^{-16}$). Our results further revealed a substantial shared proportion ($\text{Var}^2_{\text{language-shared}} = 65.78\%$), estimated from the square root of averaged correlation among cross-language pairs. The language-specific component ($\text{Var}^2_{\text{language-specific}} = 4.20\%$) was comparatively small, calculated as the difference between the square root of averaged within-language correlations and the shared component. The remaining variance ($\text{Var}^2_{\text{individual}} = 30.02\%$) was attributed to individual differences and sampling error.

Study 3: Validation of commonalities (neural activation data)

Methods: To assess the presence of language-specific components in neural representations, we conducted a comparative analysis of within-language and cross-language neural similarities similar to Study 2. Inter-subject correlation matrices were constructed from individual t-value maps for the previously identified ROIs. We employed a two-sided Wilcoxon signed-rank test to compare neural similarities between within-language subject pairs and cross-language subject pairs. Due to the shallow sampling approach used in the dataset (one or two speakers per language), precise estimation of the variance in commonalities and variations was not feasible.

Results: To assess the commonality of neural activity in our ROIs, we compared cross-language similarity (N = 7,228 pairs) with within-language similarity (N = 82 pairs). Results showed no significant differences between within-language and between-language pair similarities in the right ATL region: $M_{within}$ = 0.21, $M_{between}$ = 0.20, Wilcoxon test W = 284356, P = 0.539, suggesting a relatively common nature of neural representations in these regions. This finding aligns with previous analyses of the whole language network (Malik-Moraleda et al., 2022).

**S5 Environmental prediction results of the right ATL in nonlinguistic tasks**

To determine whether the observed climate effects on neural patterns were specific to language processing or reflected more general environmental influences on brain activity, we conducted control analyses using neural data from nonlinguistic cognitive tasks. We examined the same participants' neural responses in the right anterior temporal lobe (r-ATL) during two control tasks: Spatial Working Memory (Hard vs Easy contrast) and Math (Hard vs Easy contrast).

For each control task, we constructed neural representational dissimilarity matrices (RDMs) following the same procedures used for the language task and tested whether environmental variables (climate, cultural, geographic, and linguistic distances) predicted neural pattern dissimilarities using linear mixed-effects regression models with random intercepts for language families.

For the Spatial Working Memory task (Figure S7a), we found that cultural distance significantly predicted neural pattern dissimilarities (β = 0.13, 95% CI [0.02, 0.24], P = 0.016), while climate distance showed no significant effect (β = 0.06, 95% CI [-0.05, 0.16], P = 0.213). Geographic distance (β = -0.03, 95% CI [-0.10, 0.05], P = 0.501) and linguistic distance (β = -0.003, 95% CI [-0.07, 0.07], P = 0.925) also showed no significant effects. For the Math task (Figure S7b), none of the environmental variables significantly predicted neural pattern dissimilarities (all Ps > 0.05).

These results stand in contrast to our findings from the language task, where climate distance was the only significant predictor of neural pattern dissimilarities in the r-ATL. The dissociation between language and nonlinguistic tasks suggests that the climate effects observed during language processing are specific to linguistic mechanisms rather than reflecting general effects of environment on neural activity.

**S6 Multiple proofreading translation validation**

First, professional translation from English to the target languages (Version A) was conducted. Version A was then back-translated into English using both ChatGPT (https://chat.openai.com) and deep-L Translate (https://www.deepl.com) to allow for direct comparison with the original version. A skilled bilingual individual proofread Version A and addressed any inconsistencies identified during the back translation, leading to the development of Version B. Version B was then submitted to a second professional translator for review, focusing on resolving previously identified corrections. This iterative process was repeated 1 to 3 times for each language until a consensus was reached on the final translation.

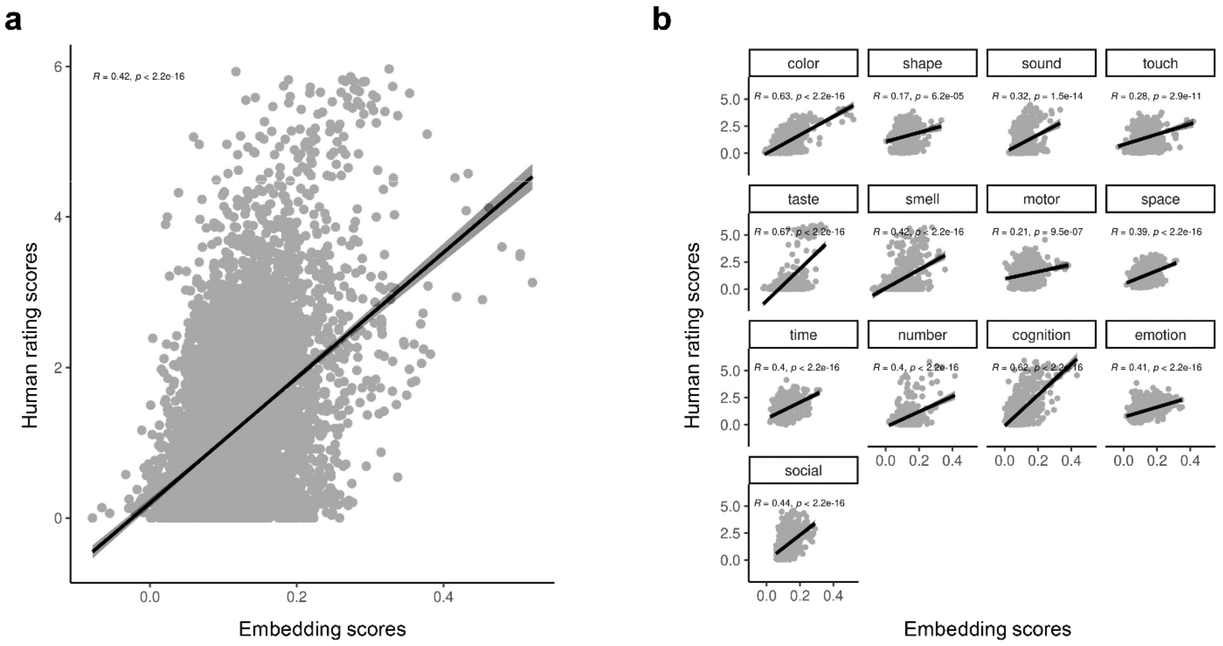

**Figure S1. Validation of the language model as a measurement tool for probing neurocognitive semantic knowledge.** We used a set of 535 English words and their human semantic ratings from (Binder et al., 2016) to examine the correlations between these ratings and word-dimension relations derived from the language model. **a**. The scatter plot of the 13-dimensional relations between human and language models, which demonstrate a moderate correlation (Pearson r = 0.42, P < 0.001). **b**. Scatter plots of word-dimension relations for each semantic dimension. As some of our dimensions correspond to several ratings in (Binder et al., 2016), for these dimensions, the human rating data were first averaged and then correlated with our data derived from language models. The detailed correspondence between our dimensions and the averaged dimensions in (Binder et al., 2016) is shown as follows: shape-Shape, Large, Small, Pattern, and Complexity; color-Color, Dark, and Bright; touch-Touch, Temperature, Texture, Weight, and Pain; sound-Sound, Audition, Loud, Low, High, and Music; taste-Taste; smell-Smell; motor-Head, Upper limb, and Lower limb; space-Landmark, Path, Scene, Near, Toward, and Away; number-Number; time-Time, Duration, Long, Short, Caused, and Consequential; social-Social, Human, Communication, and Self; cognition-Cognition; emotion-Pleasant, Unpleasant, Happy, Sad, Angry, Disgusted, Fearful, and Surprised.

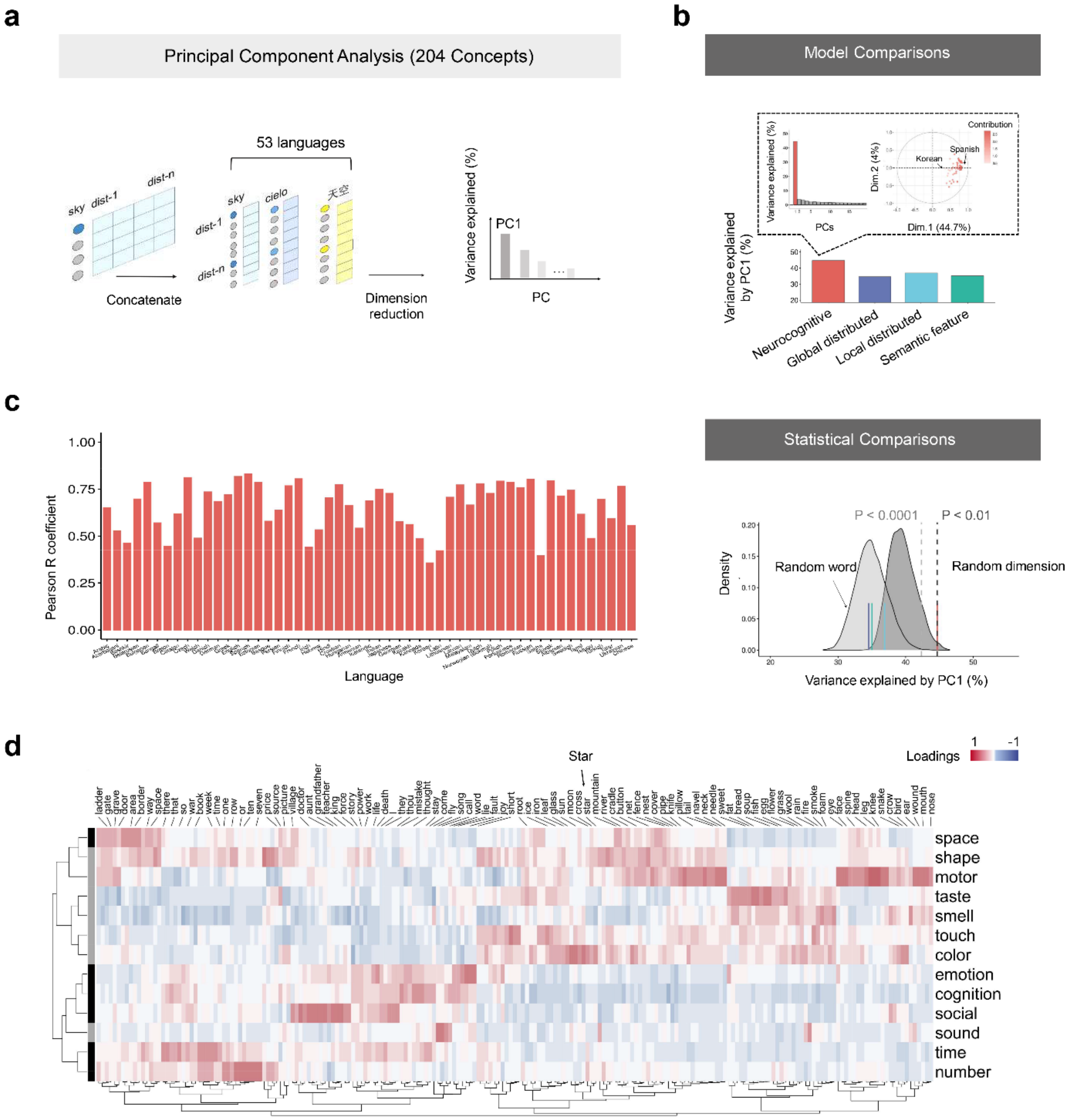

**Figure S2**. **Principal component results for 204 concepts shared across 53 languages. a.** Scheme of analyzing commonalities using principal component analysis. **b**. Principal component results. Upper panel: the bar plot shows the variance explained by PC1 for the four semantic models, with the neurocognitive model exhibiting the highest variance explained. Above the bar plot is the scree plot and the distribution of languages along the first two PCs for the neurocognitive model. Lower panel: comparison of PC1-explained variance between the four semantic models and the two random control models. The neurocognitive model reached a significance of $P < 0.0001$ in the random word distribution (light gray area) and of $P = 0.01$ in the random dimension distribution (dark gray area), while other models did not approach significance in either random control distribution. **c.** Correlation plots for all 53 languages with the first principal component of the neurocognitive semantic structure. Positive correlations suggest that this component is present across all languages. **d.** Visualization of the PC1 neurocognitive semantic space: Shared loading patterns of 204 concepts on the 13 neurocognitive dimensions across 53 languages (red: high loading; blue: low loading). Dendrograms show the hierarchical clustering of concepts (columns) and of dimensions (rows; light gray: sensory-motor; black: social-cognitive) by correlation distances.

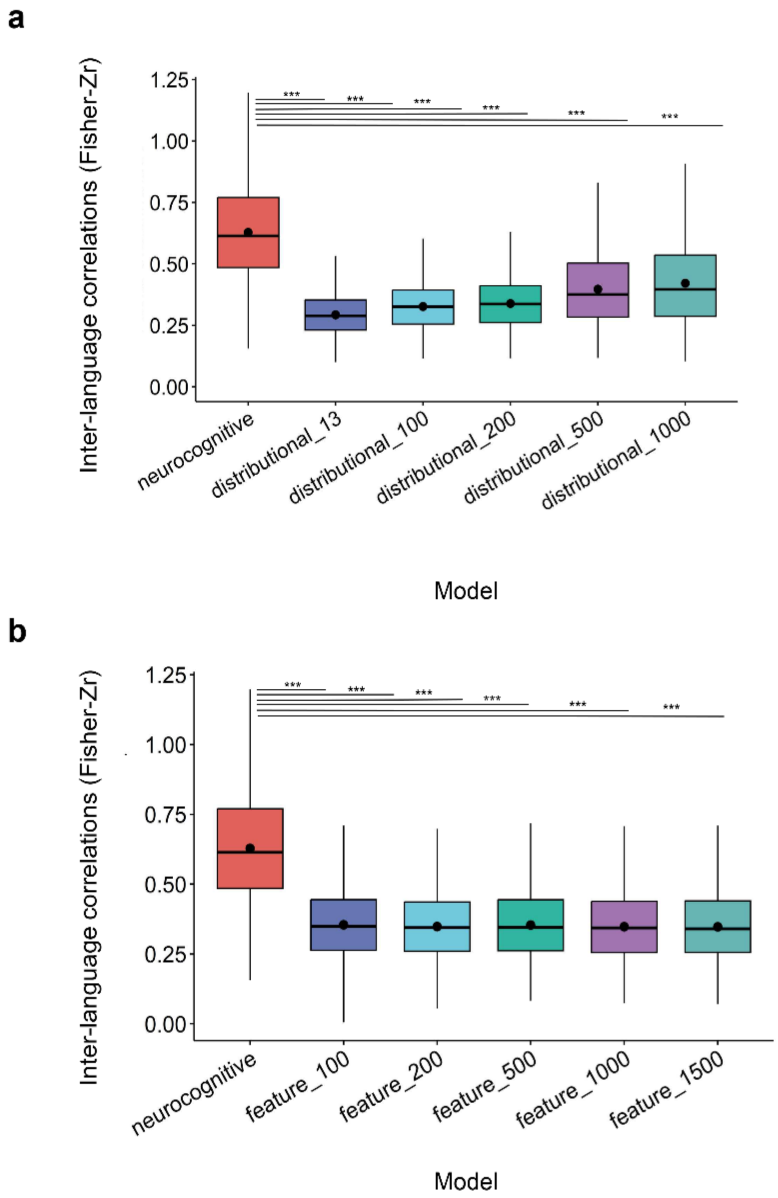

**Figure S3. Validation of inter-language correlation results of distributional (local) and feature models with varying numbers of anchor words.** In the main text, we compared the inter-language correlations of neurocognitive models with 100 anchor words in the distributional (local) model and in the feature model. Here, we validated the results using different numbers of anchor words. **a.** Distributional (local) model results. Concepts from the neurocognitive model exhibited stronger alignment than the distributional (local) models across anchor word counts ranging from 13 to 1000. **b**. Feature model results. Concepts from the neurocognitive model demonstrated superior alignment compared to feature models across feature word counts spanning 100 to 1000. Significance levels: ***, P < 0.001.

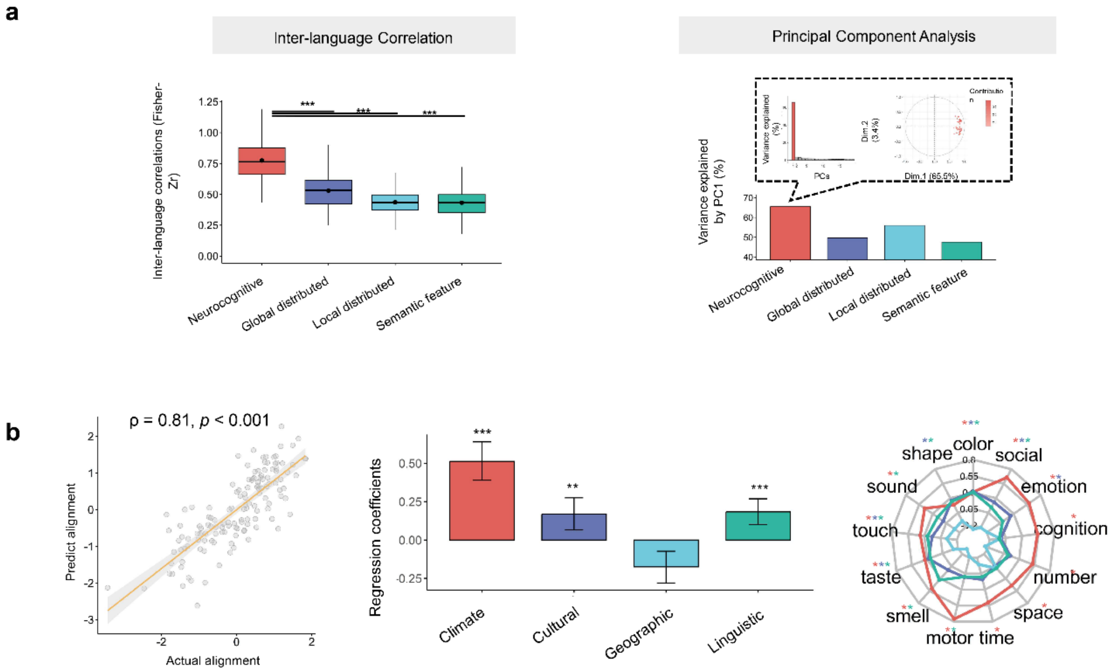

**Figure S4. Validation of the main results using independent language models trained on the combination of Wikipedia and the Open Subtitle database.** The figure represents validation analyses using word embedding models available for 35 languages (Van Paridon & Thompson, 2021), replicating main results on semantic commonalities and variation. **a.** Commonality analyses. Left panel: Inter-language correlation results. Fisher-Z transformed Pearson correlation values were calculated for each language pair for each semantic model. The neurocognitive semantic model shows greater alignment than the distributional or feature-defined semantic models. Error bars indicate 95% confidence intervals. Right panel: PCA results for 35 languages and 354 concepts. Variance explained by the first component (PC1) indicates higher commonalities in neurocognitive-based representation (65.62%) compared to other types of semantic structures (distributional--global: 45.28%; distributional-local: 55.94%; feature-based: 47.64%). Above the bar plot is the scree plot and the distribution of languages along the first two PCs for the neurocognitive model. **b**. Left panel: Scatter plot showing the relationship between actual and predicted semantic alignment values based on environmental variables. Middle panel: Bar plot showing standardized regression coefficients for each environmental variable using linear mixed models with a crossed random-effects structure that nested language pairs within language families. Climate effects were replicated, with cultural and linguistic distance also showing significant effects on semantic variations. Right panel: Radar plot displaying standardized beta coefficients from regression models for each of the 13 neurocognitive dimensions across environmental variables. Significance levels: ***, P < 0.001, **, P < 0.01, *, P < 0.05.

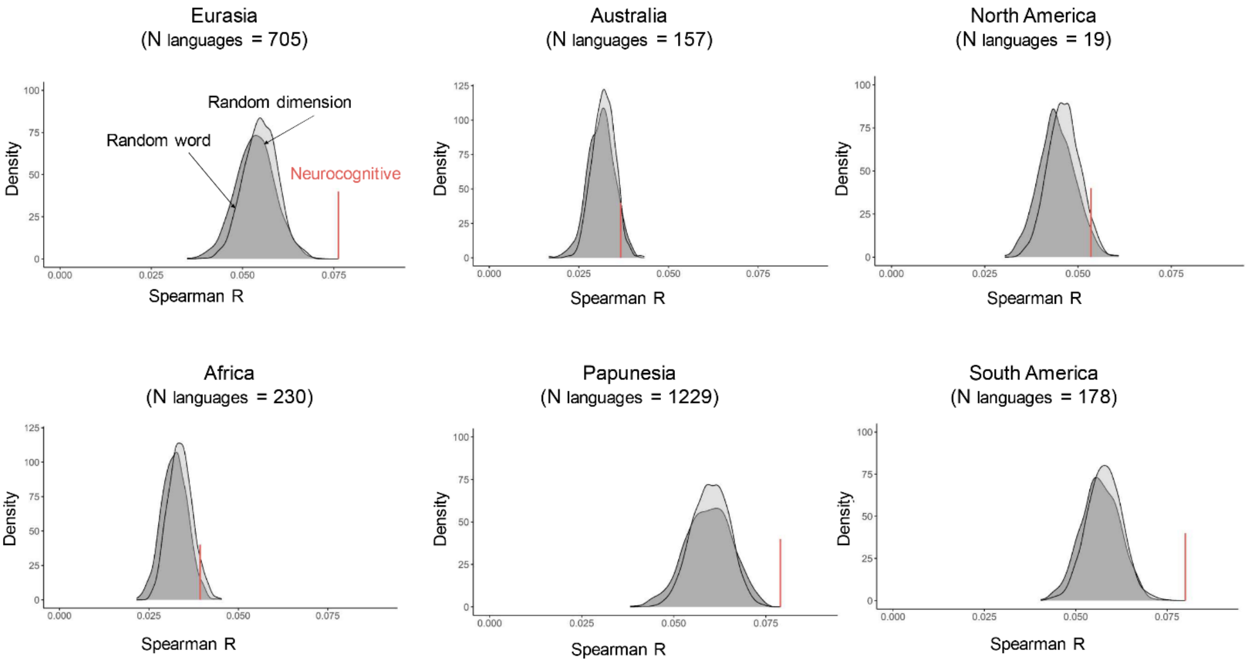

**Figure S5 Generalization of 13-dimensional neurocognitive space to language samples in different geographic region areas using the colexification network topology**. Density plots comparing association patterns between semantic distances in the original 53-language embedding-derived neurocognitive space and the colexification topological graph space. Analysis includes colexification networks from different language samples (Eurasia, Australia, North America, Africa, Papunesia, and South America). The neurocognitive model's performance is at the upper bound of the random model distributions.

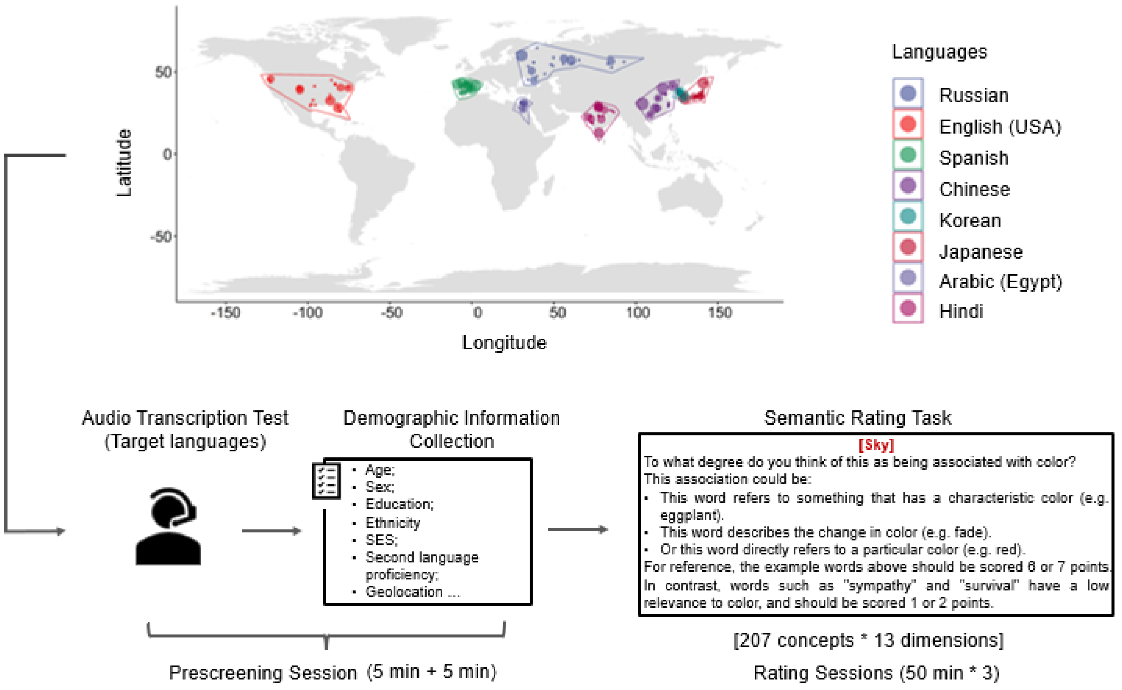

**Figure S6 Sampling and data collection (Study 2).** Diagrams outline the procedures for obtaining human ratings on 207 Swadesh concepts across 13 neurocognitive dimensions. We selected languages from Study 1, ensuring geographic, linguistic, and cultural diversity. Native speakers of 8 languages were recruited from 58 cities across 8 countries (Russia, USA, Spain, China, South Korea, Japan, Egypt, and India) via an online platform (http://www.appen.com). The participants completed a 5-minute audio transcription test in their native language and provided demographic information. After prescreening, they undertook the semantic rating task (over three sessions, each lasting approximately 50 minutes), associating concepts with each dimension.

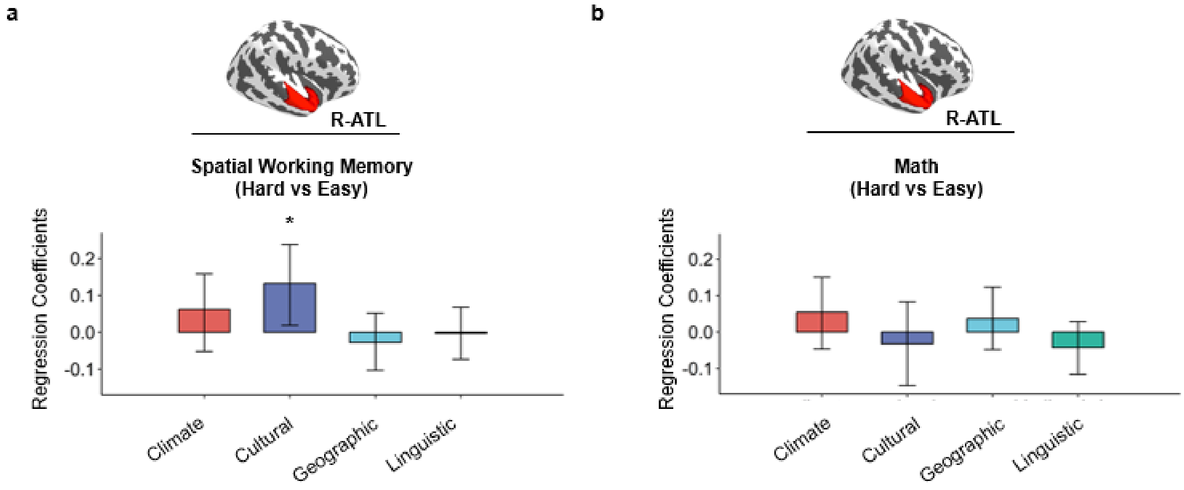

**Figure S7 Beta coefficients from linear mixed-effects regression models predicting neural pattern dissimilarities in the right anterior temporal lobe (r-ATL) during nonlinguistic tasks.** (a) Spatial Working Memory task (Hard vs Easy condition), showing significant effects only for cultural distance ($\beta = 0.13$, 95% CI [0.02, 0.24], $P = 0.016$) but not for climate distance ($\beta = 0.06$, 95% CI [-0.05, 0.16], $P = 0.213$). (b) Math task (Hard vs Easy condition), showing no significant effects for any environmental variable (all Ps > 0.05). Error bars represent 95% confidence intervals; * indicates $P < 0.05$.

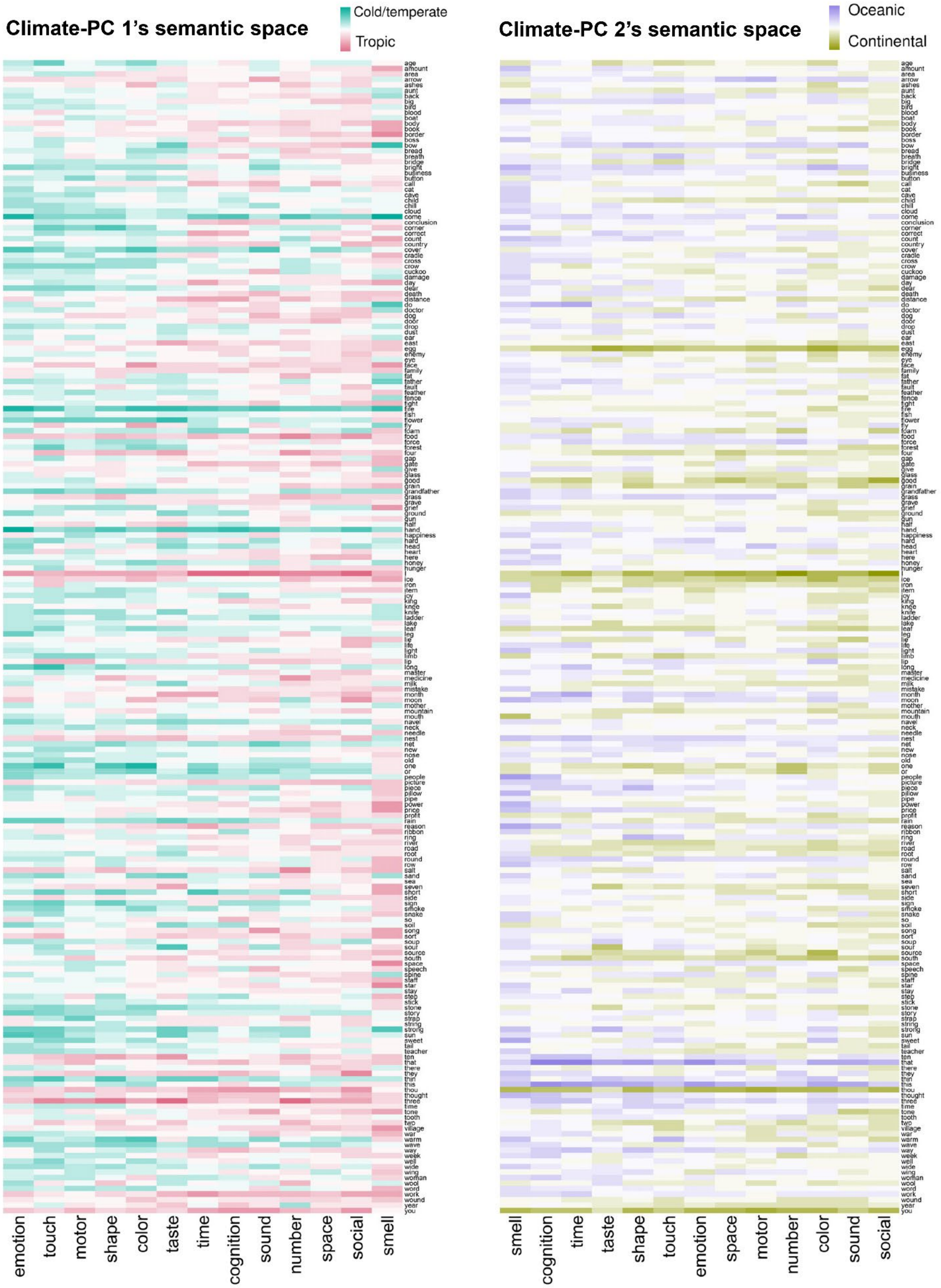

**Figure S8. Semantic profiles associated with Climate PC1 and Climate PC2 at the concept level.**

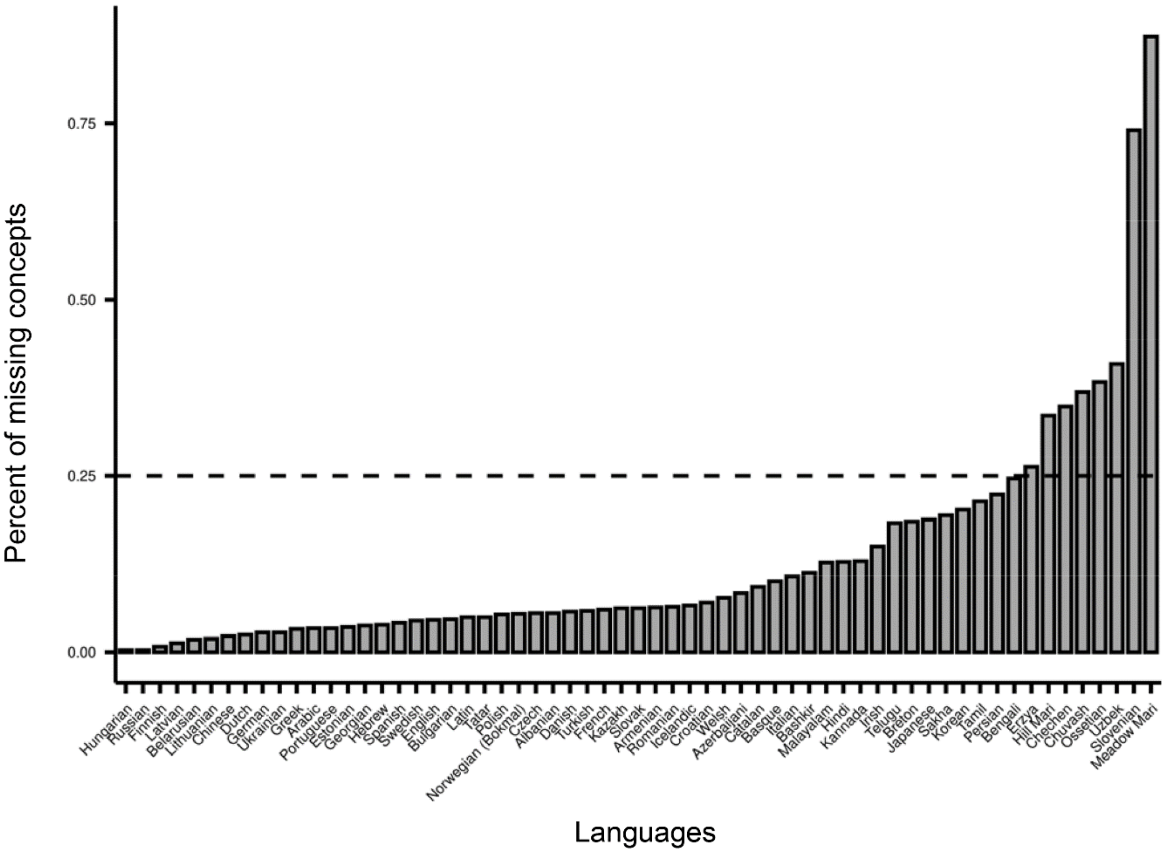

**Figure S9. Distribution of NEL concept omission in language computation models for all 61 languages.** The languages with omission percentages higher than 25% were excluded from the main analyses, including Erzya, Hill Mari, Chechen, Chuvash, Ossetian, Uzbek, Slovenian, and Meadow Mari.

**Table S1. Example (non-exhaustive) empirical references contributing to the neurocognitive dimensional framework** (gleaned from representative reviews and meta-analyses studies of the neural basis of semantic processing)**.**

| Dimensions | Task | Stimuli | Language | Contrast | Imaging modality | Year | References |
|---|---|---|---|---|---|---|---|
| Color | Word generation | pictures, words | NA | Color > Action | PET | 1995 | Martin, A., Haxby, J. V., Lalonde, F. M., Wiggs, C. L., & Ungerleider, L. G. (1995). Discrete cortical regions associated with knowledge of color and knowledge of action. Science, 270(5233), 102-105. |
| | Semantic decision | words | English | Color > Non-color (Achromatic) | fMRI | 2012 | Hsu, N. S., Frankland, S. M., & Thompson-Schill, S. L. (2012). Chromaticity of color perception and object color knowledge. Neuropsychologia, 50(2), 327-333. |
| | Semantic decision | words | Italian | Color > Action | fMRI | 2020 | Bottini, R., Ferraro, S., Nigri, A., Cuccarini, V., Bruzzone, M. G., & Collignon, O. (2020). Brain regions involved in conceptual retrieval in sighted and blind people. Journal of Cognitive Neuroscience, 32(6), 1009-1025. |
| Shape | Seman decision | words | English | Shape > (Action, Sound) | fMRI | 2003 | Noppeney, U., Friston, K. J., & Price, C. J. (2003). Effects of visual deprivation on the organization of the semantic system. Brain, 126(7), 1620-1627. |

| | | | | | | | |
|---|---|---|---|---|---|---|---|
| | Lexical decision | words | English | Visual > Sound | fMRI | 2013 | Bonner, M. F., Peelle, J. E., Cook, P. A., & Grossman, M. (2013). Heteromodal conceptual processing in the angular gyrus. NeuroImage, 71, 175-186. |
| | Semantic decision | words | Chinese | Shape RSA effects | fMRI | 2014 | Peelen, M. V., He, C., Han, Z., Caramazza, A., & Bi, Y. (2014). Nonvisual and visual object shape representations in occipitotemporal cortex: evidence from congenitally blind and sighted adults. Journal of Neuroscience, 34(1), 163-170. |
| Sound | Semantic decision | words | English | Sound > (Taste, Touch, Color) | fMRI | 2006 | Goldberg, R. F., Perfetti, C. A., & Schneider, W. (2006). Perceptual knowledge retrieval activates sensory brain regions. Journal of Neuroscience, 26(18), 4917-4921. |
| | Lexical decision | words | English | Sound > Visual | fMRI | 2013 | Bonner, M. F., Peelle, J. E., Cook, P. A., & Grossman, M. (2013). Heteromodal conceptual processing in the angular gyrus. NeuroImage, 71, 175-186. |
| | Lexical decision | words | German | Sound > Non-sound | fMRI | 2008 | Kiefer, M., Sim, E. J., Herrnberger, B., Grothe, J., & Hoenig, K. (2008). The sound of concepts: Four markers for a link between auditory and conceptual brain systems. Journal of Neuroscience, 28(47), 12224-12230. |

| | | | | | | | |
|---|---|---|---|---|---|---|---|
| Touch | Semantic decision | words | English | Touch > (Sound, Taste, Color) | fMRI | 2006 | Goldberg, R. F., Perfetti, C. A., & Schneider, W. (2006). Perceptual knowledge retrieval activates sensory brain regions. Journal of Neuroscience, 26(18), 4917-4921. |
| | Mental imagery | pictures | NA | Hand > Body | fMRI | 2016 | Perruchoud, D., Michels, L., Piccirelli, M., Gassert, R., & Ionta, S. (2016). Differential neural encoding of sensorimotor and visual body representations. Scientific Reports, 6(1), 37259. |
| Taste | Semantic decision | words | English | Taste > (Sound, Touch, Color) | fMRI | 2006 | Goldberg, R. F., Perfetti, C. A., & Schneider, W. (2006). Perceptual knowledge retrieval activates sensory brain regions. Journal of Neuroscience, 26(18), 4917-4921. |
| | Passive reading | words | Spanish | Taste > Non-taste | fMRI | 2012 | Barros-Loscertales, A., González, J., Pulvermüller, F., Ventura-Campos, N., Bustamante, J. C., Costumero, V., ... & Ávila, C. (2012). Reading salt activates gustatory brain regions: fMRI evidence for semantic grounding in a novel sensory modality. Cerebral Cortex, 22(11), 2554-2563. |
| | Semantic decision | words | Italian | Food > (People, Place) | fMRI | 2020 | Fairhall, S. L. (2020). Cross recruitment of domain-selective cortical representations enables flexible semantic knowledge. Journal of Neuroscience, 40(15), 3096-3103. |

| | | | | | | | |
|---|---|---|---|---|---|---|---|
| Smell | Passive reading | words | Spanish | Smell > Non-smell | fMRI | 2006 | González, J., Barros-Loscertales, A., Pulvermüller, F., Meseguer, V., Sanjuán, A., Belloch, V., & Ávila, C. (2006). Reading cinnamon activates olfactory brain regions. NeuroImage, 32(2), 906-912. |
| | Semantic decision | sentences | German | Smell > Non-smell | fMRI | 2018 | Pomp, J., Bestgen, A. K., Schulze, P., Müller, C. J., Citron, F. M., Suchan, B., & Kuchinke, L. (2018). Lexical olfaction recruits olfactory orbitofrontal cortex in metaphorical and literal contexts. Brain and Language, 179, 11-21. |
| Motor | Passive reading | words | English | Action > # | fMRI | 2004 | Hauk, O., Johnsrude, I., & Pulvermüller, F. (2004). Somatotopic representation of action words in human motor and premotor cortex. Neuron, 41(2), 301-307. |
| | Word generation | pictures, words | NA | Action > Color | PET | 1995 | Martin, A., Haxby, J. V., Lalonde, F. M., Wiggs, C. L., & Ungerleider, L. G. (1995). Discrete cortical regions associated with knowledge of color and knowledge of action. Science, 270(5233), 102-105. |

| | | | | | | | |
|---|---|---|---|---|---|---|---|
| | Semantic decision | passage | English | Action > (Percept, Emotion) | fMRI | 2014 | Chow, H. M., Mar, R. A., Xu, Y., Liu, S., Wagage, S., & Braun, A. R. (2014). Embodied comprehension of stories: interactions between language regions and modality-specific neural systems. Journal of Cognitive Neuroscience, 26(2), 279-295. |
| Time | Sentence judgment | sentences | English | Time > Non-time | fMRI | 2016 | Lai, V. T., & Desai, R. H. (2016). The grounding of temporal metaphors. Cortex, 76, 43-50. |
| Space | Semantic decision | words | NA | Space > Color | PET | 1998 | Mummery, C. J., Patterson, K., Hodges, J. R., & Price, C. J. (1998). Functional neuroanatomy of the semantic system: divisible by what? Journal of Cognitive Neuroscience, 10(6), 766-777. |
| | Naming | pictures | English | Space > Object | PET | 2001 | Damasio, H., Grabowski, T. J., Tranel, D., Ponto, L. L., Hichwa, R. D., & Damasio, A. R. (2001). Neural correlates of naming actions and of naming spatial relations. NeuroImage, 13(6), 1053-1064. |

| | | | | | | | |
|---|---|---|---|---|---|---|---|
| | Semantic decision | pictures, words | Italian | Place > People | fMRI | 2014 | Fairhall, S. L., Anzellotti, S., Ubaldi, S., & Caramazza, A. (2014). Person-and place-selective neural substrates for entity-specific semantic access. Cerebral Cortex, 24(7), 1687-1696. |
| Number | Semantic decision | words | NA | Number > Animal | PET | 2005 | Thioux, M., Pesenti, M., Costes, N., De Volder, A., & Seron, X. (2005). Task-independent semantic activation for numbers and animals. Cognitive Brain Research, 24(2), 284-290. |
| | Sentence comprehension | sentences | English | Number > Non-number | fMRI | 2005 | McMillan, C. T., Clark, R., Moore, P., Devita, C., & Grossman, M. (2005). Neural basis for generalized quantifier comprehension. Neuropsychologia, 43(12), 1729-1737. |
| Emotion | Perceptual decision | words | NA | Threat > Neutral | PET | 1999 | Isenberg, N., Silbersweig, D., Engelien, A., Emmerich, S., Malavade, K., Beattie, B., ... & Stern, E. (1999). Linguistic threat activates the human amygdala. Proceedings of the National Academy of Sciences, 96(18), 10456-10459. |
| | Lexical decision | words | English | Negative > Neural | fMRI | 2006 | Nakic, M., Smith, B. W., Busis, S., Vythilingam, M., & Blair, R. J. R. (2006). The impact of affect and frequency on lexical decision: the role of the amygdala and inferior frontal cortex. NeuroImage, 31(4), 1752-1761. |

| | | | | | | | |
|---|---|---|---|---|---|---|---|
| | Semantic decision | words | English | Emotion > Non-emotion | fMRI | 2019 | Wang, X., Wang, B., & Bi, Y. (2019). Close yet independent: Dissociation of social from valence and abstract semantic dimensions in the left anterior temporal lobe. Human Brain Mapping, 40(16), 4759-4776. |
| Cognition | Sentence comprehension | sentences | English | Mental > Action | fMRI | 2012 | Kana, R. K., Blum, E. R., Ladden, S. L., & Ver Hoef, L. W. (2012). “How to do things with Words”: Role of motor cortex in semantic representation of action words. Neuropsychologia, 50(14), 3403-3409. |
| | Mental imagery | words | Italian | Mental > Physical | fMRI | 2014 | Tomasino, B., Fabbro, F., & Brambilla, P. (2014). How do conceptual representations interact with processing demands: An fMRI study on action- and abstract-related words. Brain Research, 1591, 38-52. |
| Social | Story comprehension | passages | English | Social > Physical | PET | 1995 | Fletcher, P. C., Happe, F., Frith, U., Baker, S. C., Dolan, R. J., Frackowiak, R. S., & Frith, C. D. (1995). Other minds in the brain: a functional imaging study of “theory of mind” in story comprehension. Cognition, 57(2), 109-128. |
| | Semantic decision | words | English | Social > Non-social | fMRI | 2012 | Contreras, J. M., Banaji, M. R., & Mitchell, J. P. (2012). Dissociable neural correlates of stereotypes and other forms of semantic knowledge. Social Cognitive and Affective Neuroscience, 7(7), 764-770. |

| | | | | | | |
|---|---|---|---|---|---|---|
| | Semantic decision | words | Chinese | Social > Non-social | fMRI | 2019 | Lin, N., Xu, Y., Wang, X., Yang, H., Du, M., Hua, H., & Li, X. (2019). Coin, telephone, and handcuffs: Neural correlates of social knowledge of inanimate objects. Neuropsychologia, 133, 107187. |

**Table S2 Language samples included across three studies**

| Language | ISO3* | Language family | Study |
|---|---|---|---|
| Afrikaans | afr | Indo-European | Study 3 |
| Albanian | sqi | Indo-European | Study 1 |
| Arabic | arb | Afro-Asiatic | Study 1, Study 2, Study 3 |
| Armenian | hye | Indo-European | Study 1, Study 3 |
| Azerbaijani | azj | Turkic | Study 1 |
| Bashkir | bak | Turkic | Study 1 |
| Basque | eus | Basque | Study 1, Study 3 |
| Belarusian | bel | Indo-European | Study 1, Study 3 |
| Bengali | ben | Indo-European | Study 1 |
| Breton | bre | Indo-European | Study 1 |
| Bulgarian | bul | Indo-European | Study 1, Study 3 |
| Catalan | cat | Indo-European | Study 1, Study 3 |
| Chinese | cmn | Sino-Tibetan | Study 1, Study 2, Study 3 |
| Croatian | hrv | Indo-European | Study 1, Study 3 |
| Czech | ces | Indo-European | Study 1, Study 3 |
| Danish | dan | Indo-European | Study 1, Study 3 |
| Dutch | nld | Indo-European | Study 1, Study 3 |
| English | eng | Indo-European | Study 1, Study 2, Study 3 |
| Estonian | ekk | Uralic | Study 1 |
| Farsi | pes | Indo-European | Study 3 |
| Finnish | fin | Uralic | Study 1, Study 3 |
| French | fra | Indo-European | Study 1, Study 3 |
| Georgian | kat | Kartvelian | Study 1 |
| German | deu | Indo-European | Study 1, Study 3 |
| Greek | ell | Indo-European | Study 1, Study 3 |
| Gujarati | guj | Indo-European | Study 3 |
| Hebrew | heb | Afro-Asiatic | Study 1, Study 3 |
| Hindi | hin | Indo-European | Study 1, Study 2, Study 3 |
| Hungarian | hun | Uralic | Study 1, Study 3 |
| Icelandic | isl | Indo-European | Study 1 |
| Irish | gle | Indo-European | Study 1, Study 3 |
| Italian | ita | Indo-European | Study 1, Study 3 |
| Japanese | jpn | Japonic | Study 1, Study 2, Study 3 |
| Kannada | kan | Dravidian | Study 1 |
| Kazakh | kaz | Turkic | Study 1 |
| Korean | kor | Koreanic | Study 1, Study 2, Study 3 |
| Latin | lat | Indo-European | Study 1 |
| Latvian | lav | Indo-European | Study 1, Study 3 |
| Lithuanian | lit | Indo-European | Study 1, Study 3 |
| Malayalam | mal | Dravidian | Study 1 |
| Marathi | mar | Indo-European | Study 3 |
| Nepali | npi | Indo-European | Study 3 |

| | | | |
|---|---|---|---|
| Norwegian (Bokmal) | nor | Indo-European | Study 1, Study 3 |
| Persian | pes | Indo-European | Study 1 |
| Polish | pol | Indo-European | Study 1, Study 3 |
| Portuguese | por | Indo-European | Study 1, Study 3 |
| Romanian | ron | Indo-European | Study 1, Study 3 |
| Russian | rus | Indo-European | Study 1, Study 2, Study 3 |
| Sakha | sah | Turkic | Study 1 |
| Slovak | slk | Indo-European | Study 1 |
| Slovene | slv | Indo-European | Study 3 |
| Spanish | spa | Indo-European | Study 1, Study 2, Study 3 |
| Swahili | swh | Niger-Congo | Study 3 |
| Swedish | swe | Indo-European | Study 1, Study 3 |
| Tagalog | tgl | Austroasiatic | Study 3 |
| Tamil | tam | Dravidian | Study 1, Study 3 |
| Tatar | tat | Turkic | Study 1 |
| Telugu | tel | Dravidian | Study 1, Study 3 |
| Turkish | tur | Turkic | Study 1, Study 3 |
| Ukrainian | ukr | Indo-European | Study 1, Study 3 |
| Vietnamese | vie | Austroasiatic | Study 3 |
| Welsh | cym | Indo-European | Study 1 |

* Notes: International Organization for Standardization (ISO) codes.

**Table S3 Anchor words used to probe the neurocognitive semantic structure**

| Anchor words | Concept ID (in NEL) | Neurocognitive dimensions |
|---|---|---|
| eye | Auge::N | motor |
| ear | Ohr::N | |
| nose | Nase::N | |
| mouth | Mund::N | |
| tooth | Zahn::N | |
| tongue | Zunge::N | |
| shoulder | Schulter::N | |
| arm | Arm::N | |
| elbow | Ellenbogen::N | |
| hand | Hand::N | |
| finger | Finger::N | |
| foot | Fuß::N | |
| leg | Bein::N | |
| body | Körper::N | |
| bright | hell::A | color |
| dark | dunkel::A | |
| black | schwarz::A | |
| white | weiß::A | |
| red | rot::A | |
| yellow | gelb::A | |
| blue | blau::A | |
| green | grün::A | |
| grey | grau::A | |
| shine | scheinen[Sonne]::V | |
| colourful | bunt::A | |
| hard | hart::A | touch |
| soft | weich::A | |
| heavy | schwer::A | |
| hot | heiß::A | |
| warm | warm::A | |
| cold | kalt::A | |
| damp | feucht::A | |
| wet | nass::A | |
| dry | trocken::A | |
| sharp | scharf::A | |
| blunt | stumpf::A | |
| sweet | süß::A | taste |
| bitter | bitter::A | |
| flavour | Geschmack::N | |
| salt | Salz::N | |
| sour | sauer::A | |
| odour | Geruch::N | smell |

| | | |
|---|---|---|
| smell | riechen::V | |
| noise | Lärm::N | sound |
| sound | Laut::N | |
| tone | Ton::N | |
| hear | vernehmen::V | |
| listen | zuhören::V | |
| calm | Ruhe::N | |
| circle | Kreis::N | shape |
| line | Linie::N | |
| size | Größe::N | |
| long | lang::A | |
| short | kurz::A | |
| big | groß::A | |
| little | klein::A | |
| round | rund::A | |
| space | Platz::N | space |
| place | Ort::N | |
| side | Seite::N | |
| middle | Mitte::N | |
| edge | Kante::N | |
| near | nah::A | |
| far | fern::A | |
| left | linker::A | |
| right | rechter::A | |
| down | hinab::ADV | |
| up | hinauf::ADV | |
| ahead | geradeaus::ADV | |
| behind | hinter::PRP | |
| end | Schluss::N | time |
| time | Zeit::N | |
| old | alt::A | |
| new | neu::A | |
| long | lange::ADV | |
| suddenly | plötzlich::ADV | |
| instantly | sofort::ADV | |
| later | später::ADV | |
| now | jetzt::ADV | |
| soon | bald::ADV | |
| before | vorher::ADV | |
| afterwards | danach::ADV | |
| end | enden::V | |
| begin | beginnen::V | |
| full | voll::A | number |
| empty | leer::A | |

| | | |
|---|---|---|
| half | Hälfte::N | |
| one | eins::NUM | |
| two | zwei::NUM | |
| three | drei::NUM | |
| amount | Menge::N | |
| memory | Gedächtnis::N | cognition |
| mind | Verstand::N | |
| think | denken::V | |
| believe | glauben::V | |
| understand | verstehen::V | |
| know | wissen::V | |
| remember | sich erinnern an::V | |
| forget | vergessen::V | |
| hence | von hier::ADV | |
| reason | Grund::N | |
| good | gut::A | emotion |
| bad | schlecht::A | |
| want | wollen::V | |
| love | lieben::V | |
| happiness | Glück::N | |
| grief | Kummer::N | |
| afraid | fürchten::V | |
| annoy | ärgern::V | |
| sad | traurig::A | |
| joy | Freude::N | |
| desire | Lust::N | |
| merry | lustig::A | |
| human | Mensch::N | social |
| man | Mann::N | |
| woman | Frau::N | |
| child | Kind::N | |
| family | Familie::N | |
| parents | Eltern::N | |
| people | Leute::N | |
| nation | Volk::N | |
| help | Hilfe::N | |
| friend | Freund::N | |
| enemy | Feind::N | |
| I | ich::PRN | |
| we | wir::PRN | |
| you | ihr::PRN | |
| they | sie::PRN | |

**Table S4 Correlational matrix between environmental variables and semantic variations (Study 1)**

| | Climate | Linguistic history | Geography | Culture | Semantic |
|---|---|---|---|---|---|
| Climate | - | - | - | - | - |
| Linguistic history | 0.48*** | - | - | - | - |
| Geography | 0.79*** | 0.67*** | - | - | - |
| Culture | 0.73*** | 0.37*** | 0.52*** | - | - |
| Semantic | 0.53*** | 0.39*** | 0.45*** | 0.49*** | - |

Note. ***, $P < 0.001$

**Table S5 Geographic sampling of participants in Study 2**

| Country | City | Number of participants |
|---|---|---|
| Russia | Volga Federal District | 6 |
| | South Federal District | 5 |
| | Central Federal District | 5 |
| | Urals Federal District | 5 |
| | Siberian Federal District | 5 |
| | North West Federal District | 5 |
| USA | Pennsylvania | 5 |
| | Florida | 5 |
| | Colorado | 5 |
| | Alabama | 4 |
| | Michigan | 4 |
| | Texas | 4 |
| | Oregon | 4 |
| | Tennessee | 1 |
| | Nebraska | 1 |
| Spain | Andalusia | 5 |
| | Valencian Community | 5 |
| | Community of Madria | 4 |
| | Catalonia | 4 |
| | Asturias | 3 |
| | Basque Country | 3 |
| | Extremadura | 3 |
| | Galicia | 3 |
| China | Sichuan | 5 |
| | Hunan | 5 |
| | Zhejiang | 4 |
| | Henan | 4 |
| | Shandong | 4 |
| | Liaoning | 4 |
| | Beijing | 4 |
| | Guangxi | 3 |
| South Korea | Busan | 6 |
| | Seoul | 5 |
| | Incheon | 5 |
| | Gyeonggi-do | 5 |
| | Gyeongsangnam-do | 4 |
| | Daejeon | 3 |
| | Daegu | 3 |
| Japan | Tokyo | 5 |
| | Hokoaido | 5 |
| | Aichi | 5 |
| | Kanagawa | 5 |

| | | |
|---|---|---|
| | Osaka | 5 |
| | Fukuoka | 5 |
| | Miyagi Prefecture | 1 |
| Egypt | Cairo | 8 |
| | Dakahlia | 6 |
| | Alexandria | 5 |
| | Ismailia | 5 |
| | Giza | 5 |
| | Asyut | 5 |
| India | Delhi | 6 |
| | Maharashtra | 5 |
| | Gujarat | 5 |
| | Madhya Pradesh | 4 |
| | Uttar Pradesh | 4 |
| | Bihar | 3 |
| | Karnataka | 3 |

**Table S6 Word lists of Swadesh 207 concepts in 8 languages**

| No. | English (en) | Chinese (cn) | Arabic (ar) | Hindi (hi) | Japanese (ja) | Korean (ko) | Russian (ru) | Spanish (es) |
|---|---|---|---|---|---|---|---|---|
| 1 | I | 我 | أنا | मैं | 私 | 나 | я | yo |
| 2 | you (singular) | 你 | أنت | तुम | あなた | 너 | ты | tú |
| 3 | he | 他 | هو | वह | 彼 | 그 | он | él |
| 4 | we | 我们 | نحن | हम | 私たち | 우리 | мы | nosotros |
| 5 | you (plural) | 你们 | أنتم | आप | あなた達 | 너희들 | вы | vosotros |
| 6 | they | 他们 | هم | वे | 彼ら | 그들 | они | ellos |
| 7 | this | 这个 | هذا | यह | この | 이- | это | este |
| 8 | that | 那个 | ذلك | वह | あの | 그- | то | aquel |
| 9 | here | 这里 | هنا | यहाँ | ここ | 이곳 | тут | aquí |
| 10 | there | 那里 | هناك | वहाँ | あそこ | 그곳 | там | allí |
| 11 | who | 谁 | من | कौन | 誰 | 누구 | кто | quien |
| 12 | what | 什么 | ما | क्या | 何 | 무엇 | что | que |
| 13 | where | 哪里 | أين | कहाँ | どこ | 어디 | где | donde |
| 14 | when | 什么时候 | متى | कब | いつ | 언제 | когда | cuando |
| 15 | how | 怎么 | كيف | कैसे | どう | 어떻게 | как | como |
| 16 | not | 不是 | لا | नहीं | 無い | -아니다 | не | no |
| 17 | all | 全部 | كل | सभी | すべて | 모든 | всё (совокупность чего-либо) | todos |
| 18 | many | 多 | كثير | बहुत | 多い | 많은 | много | muchos |
| 19 | some | 一些 | بعض | कुछ | 幾つか | 어떤 | несколько | algunos |
| 20 | few | 少 | قليل | थोड़े | 少し | 조금 | мало | poco |
| 21 | other | 另外的 | أخرى | दूसरा | 他 | 다른 | другой | otro |
| 22 | one | 一 | واحد | एक | 一 | 하나 | один | uno |
| 23 | two | 二 | اثنان | दो | 二 | 둘 | два | dos |
| 24 | three | 三 | ثلاثة | तीन | 三 | 셋 | три | tres |

| | | | | | | | | |
|---|---|---|---|---|---|---|---|---|
| 25 | four | 四 | أربعة | चार | 四 | 넷 | четыре | cuatro |
| 26 | five | 五 | خمسة | पाँच | 五 | 다섯 | пять | cinco |
| 27 | big | 大 | كبير | बड़ा | 大きい | 크다 | большой (обладающий немаленьким размером) | grande |
| 28 | long | 长 | طويل | लम्बा | 長い | 길다 | длинный (большой по протяженности или длительности) | largo |
| 29 | wide | 宽 | عريض | चौड़ा | 広い | 넓다 | широкий | ancho |
| 30 | thick | 厚 | سميك | मोटा | 厚い | 두껍다 | толстый | grueso |
| 31 | heavy | 重 | ثقيل | भारी | 重い | 무겁다 | тяжелый (обладающий большим весом) | pesado |
| 32 | small | 小 | صغير | छोटा | 小さい | 작다 | маленький | pequeño |
| 33 | short | 短 | قصير | लघु | 短い | 짧다 | короткий | corto |
| 34 | narrow | 窄 | ضيق | तंग | 狭い | 좁다 | узкий | estrecho |
| 35 | thin | 细 | رفيع | पतला | 細い | 얇다 | тонкий | delgado |
| 36 | woman | 女人 | امرأة | औरत | 女 | 여자 | женщина | mujer |
| 37 | man (as adult male) | 男人 | رجل | पुरुष | 男 | 남자 | мужчина | hombre |
| 38 | man (as human being) | 人 | إنسان | इंसान | 人 | 사람 | человек | humano |
| 39 | child | 儿童 | طفل | बच्चा | 子供 | 어린이 | ребенок | niño |
| 40 | wife | 妻子 | زوجة | पत्नी | 妻 | 아내 | жена | esposa |
| 41 | husband | 丈夫 | زوج | पति | 夫 | 남편 | муж | esposo |
| 42 | mother | 妈妈 | أم | माँ | 母 | 어머니 | мать | madre |
| 43 | father | 爸爸 | أب | पिता | 父 | 아버지 | отец | padre |
| 44 | animal | 动物 | حيوان | जानवर | 動物 | 동물 | животное | animal |

| | | | | | | | | |
|---|---|---|---|---|---|---|---|---|
| 45 | fish | 鱼 | سمك | मछली | 魚 | 물고기 | рыба | pez |
| 46 | bird | 鸟 | طائر | पक्षी | 鳥 | 새 | птица | pájaro |
| 47 | dog | 狗 | كلب | कुत्ता | 犬 | 개 | собака | perro |
| 48 | louse | 虱子 | قملة | जूँ | シラミ | 이 (곤충) | вошь | piojo |
| 49 | snake | 蛇 | أفعى | साँप | 蛇 | 뱀 | змея | serpiente |
| 50 | worm | 蚯蚓 | دودة | कृमि | 虫 | 벌레 | червь | gusano |
| 51 | tree | 树 | شجرة | पेड़ | 木 | 나무 | дерево | árbol |
| 52 | forest | 森林 | غابة | वन | 林 | 숲 | лес | bosque |
| 53 | branch | 树枝 | غصن | शाखा | 枝 | 나뭇가지 | ветка (часть дерева) | rama |
| 54 | fruit | 水果 | فاكهة | फल | 果物 | 과일 | фрукт (плод) | fruto |
| 55 | seed | 种子 | بذرة | बीज | 種 | 씨 | семя | semilla |
| 56 | leaf | 叶子 | ورقة | पत्ती | 葉 | 잎 | Лист (часть растения) | hoja |
| 57 | root | 根 | جذر | जड़ | 根 | 뿌리 | корень (подземная часть дерева) | raíz |
| 58 | bark (of a tree) | 树皮 | لحاء | (पेड़ की) छाल | 樹皮 | 나무껍질 | кора | corteza |
| 59 | flower | 花 | زهرة | पुष्प | 花 | 꽃 | цветок | flor |
| 60 | grass | 草 | حشيش | घास | 草 | 풀 | трава | hierba |
| 61 | rope | 绳 | حبل | रस्सी | ロープ | 밧줄 | верёвка | cuerda |
| 62 | skin | 皮肤 | جلد | त्वचा | 皮 | 피부 | кожа | piel |
| 63 | meat | 肉 | لحم | गोश्त | 肉 | 고기 | мясо | carne |
| 64 | blood | 血 | دم | रक्त | 血 | 피 | кровь | sangre |
| 65 | bone | 骨头 | عظم | हड्डी | 骨 | 뼈 | кость | hueso |
| 66 | fat (noun) | 脂肪 | دهن | वसा | 油 | 기름 | жир | grasa |
| 67 | egg | 蛋 | بيضة | अंडा | 卵 | 알 | яйцо | huevo |
| 68 | horn | 角 | قرن | सींग; (भोंपू) | 角 | 뿔 | рог | cuerno |

| | | | | | | | | |
|---|---|---|---|---|---|---|---|---|
| 69 | tail | 尾巴 | ذيل | पूँछ | 尻尾 | 꼬리 | хвост | cola |
| 70 | feather | 羽毛 | ريشة | मुलायम पंख | 羽 | 깃털 | перо | pluma |
| 71 | hair | 头发 | شعر | केश | 毛 | 머리카락 | волос | pelo |
| 72 | head | 头 | رأس | सिर | 頭 | 머리 | голова | cabeza |
| 73 | ear | 耳朵 | أذن | कान | 耳 | 귀 | ухо | oreja |
| 74 | eye | 眼睛 | عين | आँख | 目 | 눈 (기관) | глаз | ojo |
| 75 | nose | 鼻子 | أنف | नाक | 鼻 | 코 | нос | nariz |
| 76 | mouth | 嘴 | فم | मुँह | 口 | 입 | рот | boca |
| 77 | tooth | 牙 | سن | दाँत | 歯 | 이 (기관) | зуб | diente |
| 78 | tongue (organ) | 舌头 | لسان | जीभ | 舌 | 혀 | язык | lengua |
| 79 | fingernail | 指甲 | ظفر | नाखून | 爪 | 손톱 | ноготь | uña |
| 80 | foot | 脚 | قدم | पैर | 足 | 발 | стопа | pie |
| 81 | leg | 腿 | ساق | टांग | 脚 | 다리 | нога | pierna |
| 82 | knee | 膝盖 | ركبة | घुटना | 膝 | 무릎 | колено | rodilla |
| 83 | hand | 手 | يد | हाथ | 手 | 손 | рука | mano |
| 84 | wing | 翅膀 | جناح | पंख | 翼 | 날개 | крыло | ala |
| 85 | belly | 肚子 | بطن | पेट | 腹 | 배 | живот | barriga |
| 86 | guts | 内脏 | أمعاء | अंतड़ी | 内臓 | 내장 | кишечник | intestinos |
| 87 | neck | 脖子 | عنق | गर्दन | 首 | 목 | шея | cuello |
| 88 | back | 背 | ظهر | पीठ, (पीछे) | 背 | 등 | спина | espalda |
| 89 | breast | 胸 | صدر | छाती | 胸 | 가슴 | грудь | pecho |
| 90 | heart | 心 | قلب | दिल | 心臓 | 심장 | сердце | corazón |
| 91 | liver | 肝 | كبد | कलेजा | 肝臓 | 간 | печень | hígado |
| 92 | drink | 喝 | شرب | पीना, (पेय पदार्थ) | 飲む | 마시다 | пить | beber |
| 93 | eat | 吃 | أكل | खाना | 食べる | 먹다 | есть | comer |
| 94 | bite | 咬 | عضّ | मुँह से काटना | 噛む | 물다 | кусать | morder |
| 95 | suck | 吮吸 | امتصّ | चूसना | 吸う | 빨다 | сосать | chupar |

| | | | | | | | | |
|---|---|---|---|---|---|---|---|---|
| 96 | spit | 吐 | بصق | थूकना | 唾を吐く | 침뱉다 | плевать | escupir |
| 97 | vomit | 呕吐 | تقيأ | उल्टी करना | 吐く | 토하다 | тошнить | vomitar |
| 98 | blow | 吹 | نفخ | फूँकना | 吹く | 불다 | дуть | soplar |
| 99 | breathe | 呼吸 | تنفس | साँस लेना | 呼吸する | 숨쉬다 | дышать | respirar |
| 100 | laugh | 笑 | ضحك | हँसना | 笑う | 웃다 | смеяться | reír |
| 101 | see | 看 | رأى | देखना | 見る | 보다 | видеть | ver |
| 102 | hear | 听 | سمع | सुनना | 聞く | 듣다 | слышать | oír |
| 103 | know | 知道 | علم | जानना | 知る | 알다 | знать | saber |
| 104 | think | 思考 | افتكر | सोचना | 思う | 생각하다 | думать | pensar |
| 105 | smell | 闻 | اشتم | सूँघना | 匂う | 냄새 | нюхать | oler |
| 106 | fear | 害怕 | خاف | डरना | 恐れる | 두려움 | бояться | temer |
| 107 | sleep | 睡觉 | نام | सोना | 寝る | 자다 | спать | dormir |
| 108 | live | 活 | عاش | जीवित रहना | 生きている | 살다 | жить | vivir |
| 109 | die | 死 | مات | मरना | 死ぬ | 죽다 | умирать | morir |
| 110 | kill | 杀 | قتل | जान से मारना | 殺す | 죽이다 | убивать | matar |
| 111 | fight | 打架 | تقاتل | लड़ना | 争う | 싸우다 | драться | luchar |
| 112 | hunt | 打猎 | صاد | शिकार करना | 狩る | 사냥하다 | охотиться | cazar |
| 113 | hit | 打 | ضرب | टकराना | 叩く | 때리다 | ударить | golpear |
| 114 | cut | 切开 | قطع | कटाई करना | 切る | 베다 | резать | cortar |
| 115 | split | 分开 | شقّ | विभाजित करना | 裂く | 갈라지다 | разделить | dividir |
| 116 | stab | 扎 | طعن | भोंकना | 刺す | 찌르다 | заколоть | apuñalar |
| 117 | scratch | 挠 | خدش | खरोंचना | 掻く | 긁다 | чесать | arañar |
| 118 | dig | 挖 | حفر | खोदना | 彫る | 파다 | копать | cavar |
| 119 | swim | 游泳 | سبح | तैरना | 泳ぐ | 헤엄치다 | плавать | nadar |
| 120 | fly (verb) | 飞 | طار | उड़ना | 跳ぶ | 날다 | летать | volar |
| 121 | walk | 走 | مشى | टहलना | 歩く | 걷다 | ходить | andar |
| 122 | come | 来 | أتى | आना | 来る | 오다 | приходить | venir |

| | | | | | | | | |
|---|---|---|---|---|---|---|---|---|
| 123 | lie (as in a bed) | 躺 | رقد | लेटना | 嘘を吐く | 눕다 | лежать | acostarse |
| 124 | sit | 坐 | جلس | बैठना | 座る | 앉다 | сидеть | sentarse |
| 125 | stand | 站 | وقف | खड़े होना | 立つ | 서다 | стоять | ponerse de pie |
| 126 | turn | 转 | التفت | मुड़ना | 回る | 돌다 | поворачивать | girar |
| 127 | drop | 落(luo)下 | سقط | गिराना, (बूंद) | 落ちる（落とす） | 떨어지다 | ронять | caer |
| 128 | give | 给 | أعطى | देना | あげる | 주다 | давать | dar |
| 129 | hold | 拿 | أمسك | पकड़ना | 取る | 붙잡다 | держать | apoyar |
| 130 | squeeze | 挤压 | عصر | निचोड़ना | 搾る | 짜다 | сжимать | estrujar |
| 131 | rub | 搓 | مسح | रगड़ना | 擦る | 문지르다 | тереть | frotar |
| 132 | wash | 洗 | غسل | धोना | 洗う | 씻다 | мыть | lavar |
| 133 | wipe | 擦 | محا | पोंछना | 拭く | 닦다 | вытирать | enjugar |
| 134 | pull | 拉 | جر | खींचना | 引く | 당기다 | тянуть | tirar |
| 135 | push | 推 | دفع | धकेलना | 押す | 밀다 | толкать | empujar |
| 136 | throw | 扔 | رمى | फेंकना | 投げる | 던지다 | бросать | arrojar |
| 137 | tie (verb) | 系（动词） | ألقى | बाँधना | 結ぶ | 매다 | связывать | atar |
| 138 | sew | 缝 | خاط | सिलाई करना | 縫う | 꿰매다 | шить | coser |
| 139 | count | 数（动词） | عدّ | गिनना | 数える | 세다 | Считать （называть числа в последовательном порядке) | contar |
| 140 | say | 说 | قال | कहना | 言う | 말하다 | говорить | decir |
| 141 | sing | 唱 | غنّى | गाना | 歌う | 노래하다 | петь | cantar |
| 142 | play | 玩 | لعب | खेलना, (नाटक) | 遊ぶ（弾く） | 놀다 | играть | jugar |
| 143 | float | 漂浮 | طفا | बहाना | 浮く | 뜨다 | плыть | flotar |
| 144 | flow | 流 | تدفق | बहना | 流れる | 흐르다 | течь | correr |
| 145 | freeze | 结冰 | جمد | जमना | 凍る | 얼다 | замерзать | congelar |

| | | | | | | | | |
|---|---|---|---|---|---|---|---|---|
| 146 | swell | 膨胀 | ورم | सूजना | 膨らむ（腫れる） | 붓다 | набухать | hinchar |
| 147 | sun | 太阳 | شمس | सूरज | 太陽 | 해 | солнце | sol |
| 148 | moon | 月亮 | قمر | चंद्रमा | 月 | 달 | луна | luna |
| 149 | star | 星星 | نجم | तारा | 星 | 별 | звезда | estrella |
| 150 | water | 水 | ماء | पानी | 水 | 물 | вода | agua |
| 151 | rain | 雨 | مطر | बारिश | 雨 | 비 | дождь | lluvia |
| 152 | river | 河 | نهر | नदी | 川 | 강 | река | río |
| 153 | lake | 湖 | بحيرة | झील | 池 | 연못 | озеро | lago |
| 154 | sea | 海 | بحر | समुद्र | 海 | 바다 | море | mar |
| 155 | salt | 盐 | ملح | नमक | 塩 | 소금 | соль | sal |
| 156 | stone | 石头 | حجر | पत्थर | 岩 | 돌 | камень | piedra |
| 157 | sand | 沙子 | رمل | बालू | 砂 | 모래 | песок | arena |
| 158 | dust | 灰尘 | غبار | धूल | 埃 | 먼지 | пыль | polvo |
| 159 | earth | 土地 | أرض | पृथ्वी | 土 | 땅 | земля | tierra |
| 160 | cloud | 云 | سحاب | बादल | 雲 | 구름 | облако | nube |
| 161 | fog | 雾 | ضباب | कोहरा | 霧 | 안개 | туман | niebla |
| 162 | sky | 天空 | سماء | आकाश | 空 | 하늘 | небо | cielo |
| 163 | wind | 风 | ريح | हवा | 風 | 바람 | ветер | viento |
| 164 | snow | 雪 | ثلج | प्राकृतिक बर्फ़ | 雪 | 눈 (날씨) | снег | nieve |
| 165 | ice | 冰 | جليد | बर्फ़ | 氷 | 얼음 | лед | hielo |
| 166 | smoke | 烟 | دخان | धुआँ | 煙 | 연기 | дым | humo |
| 167 | fire | 火 | نار | आग | 火 | 불 | огонь | fuego |
| 168 | ash | 灰烬 | رماد | राख | 灰 | 재 | зола | ceniza |
| 169 | burn | 烧 | احترق | जलना | 燃える（燃やす） | 타다 | гореть | arder |
| 170 | road | 道路 | طريق | सड़क | 道 | 길 | дорога | calle |
| 171 | mountain | 山 | جبل | पहाड़ | 山 | 산 | гора | montaña |
| 172 | red | 红 | أحمر | लाल | 赤い | 빨강 | красный | rojo |
| 173 | green | 绿 | أخضر | हरा | 緑 | 초록 | зеленый | verde |
| 174 | yellow | 黄 | أصفر | पीला | 黄色 | 노랑 | желтый | amarillo |

| | | | | | | | | |
|---|---|---|---|---|---|---|---|---|
| 175 | white | 白 | أبيض | सफ़ेद | 白い | 하양 | белый | blanco |
| 176 | black | 黑 | أسود | काला | 黒い | 검정 | черный | negro |
| 177 | night | 夜晚 | ليل | रात | 夜 | 밤 | ночь | noche |
| 178 | day | 白天 | نهار | दिन | 日 | 낮 | день | día |
| 179 | year | 年 | سنة | साल | 年 | 년 | год | año |
| 180 | warm | 暖和 | دافئ | गर्म | 暖かい | 따뜻하다 | теплый | caliente |
| 181 | cold | 冷 | بارد | ठंडा | 冷たい | 차다 | холодный | frío |
| 182 | full | 满 | مليء | पूरा | 満杯 | 가득차다 | полный | lleno |
| 183 | new | 新 | جديد | नया | 新しい | 새- | новый | nuevo |
| 184 | old | 陈旧的 | قديم | पुराना | 古い | 옛- | старый | viejo |
| 185 | good | 好 | جيد | अच्छा | 良い | 좋다 | хороший | bueno |
| 186 | bad | 坏 | سيء | ख़राब | 悪い | 나쁘다 | плохой | mal |
| 187 | rot | 腐烂 | عفِن | सड़ांध | 腐る | 썩다 | гнилой | podrido |
| 188 | dirty | 脏的 | وسخ | गंदा | 汚い | 더럽다 | грязный | sucio |
| 189 | straight | 直的 | مستقيم | सीधा | 真っ直ぐ | 곧다 | прямой | derecho |
| 190 | round | 圆的 | دائري | गोल | 丸い | 둥글다 | круглый | redondo |
| 191 | sharp | 锋利 | حادّ | पैना | 鋭い | 날카롭다 | острый | afilado |
| 192 | blunt | 钝 | غيْر حادّ | कुंद | 鈍い | 무디다 | тупой | desafilado |
| 193 | smooth | 光滑 | ناعم | चिकना | 滑らか（順調な、流暢な） | 부드럽다 | гладкий | suave |
| 194 | wet | 湿的 | رطب | गीला | 濡れる | 젖다 | мокрый | mojado |
| 195 | dry | 干的 | جاف | सूखा | 乾く | 마르다 | сухой | seco |
| 196 | correct | 正确 | صحيح | सही | 正解（正す） | 맞다 | правильный | correcto |
| 197 | near | 近 | قريب | समीप | 近い | 가깝다 | близкий | cerca |
| 198 | far | 远 | بعيد | दूर | 遠い | 멀다 | далекий | distante |
| 199 | right (as opposed to left) | 右 | يَمِين | दायां , (सही) | 右 | 오른쪽 | правый | derecha |
| 200 | left | 左 | يَسار | बायां, (छोड़ा) | 左 | 왼쪽 | левый | izquierda |

| | | | | | | | | |
|---|---|---|---|---|---|---|---|---|
| 201 | at | 在 | عند | पर | で | -에서 | у (около) | a |
| 202 | in | 在…里面 | فِي | में | 中 | -안에 | в | adentro |
| 203 | with | 跟 | مع | के साथ | と | -와 | с | con |
| 204 | and | 与 | و | और | そして | 그리고 | и | y |
| 205 | if | 如果 | اذا | यदि | もし | 만약--<br>한다면 | если | si |
| 206 | because | 因为 | لأنّ | क्योंकि | だから | 때문에 | потому что | porque |
| 207 | name | 名字 | اِسْم | नाम | 名前（タイトル） | 이름 | имя | nombre |

**Table S7 Profiles of two major climate PCs on 19 biologically meaningful climate variables**

| Climate variable | Variable description | PC1 loadings | PC2 loadings |
|---|---|---|---|
| BIO1 | Annual Mean Temperature | -.97 | -.06 |
| BIO2 | Mean Diurnal Range (Mean of monthly (max temp - min temp)) | -.28 | -.81 |
| BIO3 | Isothermality (BIO2/BIO7) (* 100) | -.87 | .14 |
| BIO4 | Temperature Seasonality (standard deviation *100) | .69 | -.57 |
| BIO5 | Max Temperature of Warmest Month | -.73 | -.61 |
| BIO6 | Min Temperature of Coldest Month | -.89 | .31 |
| BIO7 | Temperature Annual Range (BIO5-BIO6) | .53 | -.74 |
| BIO8 | Mean Temperature of Wettest Quarter | -.49 | -.45 |
| BIO9 | Mean Temperature of Driest Quarter | -.84 | .09 |
| BIO10 | Mean Temperature of Warmest Quarter | .82 | .49 |
| BIO11 | Mean Temperature of Coldest Quarter | .94 | .20 |
| BIO12 | Annual Precipitation | -.48 | .64 |
| BIO13 | Precipitation of Wettest Month | -.67 | .24 |
| BIO14 | Precipitation of Driest Month | .33 | .85 |
| BIO15 | Precipitation Seasonality (Coefficient of Variation) | -.62 | -.59 |
| BIO16 | Precipitation of Wettest Quarter | -.66 | .30 |
| BIO17 | Precipitation of Driest Quarter | .26 | .89 |
| BIO18 | Precipitation of Warmest Quarter | -.14 | .34 |
| BIO19 | Precipitation of Coldest Quarter | -.09 | .81 |

#### References

Binder, J. R., Conant, L. L., Humphries, C. J., Fernandino, L., Simons, S. B., Aguilar, M., et al. (2016). Toward a brain-based componential semantic representation. *Cognitive neuropsychology, 33*(3-4), 130-174.

Chen, G., Taylor, P. A., Shin, Y.-W., Reynolds, R. C., & Cox, R. W. (2017). Untangling the relatedness among correlations, Part II: Inter-subject correlation group analysis through linear mixed-effects modeling. *NeuroImage, 147*, 825-840.

Cole, M. W., Bassett, D. S., Power, J. D., Braver, T. S., & Petersen, S. E. (2014). Intrinsic and task-evoked network architectures of the human brain. *Neuron, 83*(1), 238-251.

Malik-Moraleda, S., Ayyash, D., Gallée, J., Affourtit, J., Hoffmann, M., Mineroff, Z., et al. (2022). An investigation across 45 languages and 12 language families reveals a universal language network. *Nature Neuroscience, 25*(8), 1014-1019.

Romney, A. K., Moore, C. C., Batchelder, W. H., & Hsia, T.-L. (2000). Statistical methods for characterizing similarities and differences between semantic structures. *Proceedings of the National Academy of Sciences, 97*(1), 518-523.

Van Paridon, J., & Thompson, B. (2021). subs2vec: Word embeddings from subtitles in 55 languages. *Behavior Research Methods, 53*, 629-655.
